# Mapping genomic regions associated with temperature stress in the wheat pathogen *Zymoseptoria tritici*

**DOI:** 10.1101/2024.09.23.614455

**Authors:** Jessica Stapley, Ziming Zhong, Bruce A. McDonald

**Author notes:** Corresponding author: Jessica Stapley. co-first authors (Jessica Stapley, Ziming Zhong).

## Abstract

Climate change can alter interactions between plants and their pathogens, which could adversely affect crop production. To better understand the molecular mechanisms underlying the responses of pathogenic fungal to temperature stress, we conducted a quantitative trait loci (QTL) mapping study in the wheat pathogen *Zymoseptoria tritici* to identify genomic regions associated with colony growth and melanisation at three temperatures (10°C, 18°C, 27°C). We then identified likely candidate genes for thermal adaptation within these intervals by combining information regarding gene function, GO annotation enrichment, transcriptional profile, and results from previous genome wide association studies (GWAS) investigating responses to climate, temperature and thermal adaptation. The QTL mapping, conducted for two separate crosses involving four Swiss parents, found significant QTL uniquely associated with traits measured in high and low temperatures. These intervals contained many genes known to regulate responses to temperature stress, including heat shock proteins (HSPs) and proteins involved in the mitogen-activated protein kinase (MAPK) pathways, and were enriched for genes with a zinc ion binding GO annotation. We highlight the most promising candidate genes for thermal adaptation, including an ammonium transporter gene, a stress response factor (*Whi1*) and two MAPK pathway genes - *SSk2* and *Opy2*. Future validation work on these candidate genes could provide novel insight into the molecular mechanisms underlying temperature adaptation in this important wheat pathogen.

## Introduction

Human-induced climate change is the greatest challenge facing our planet. The warming climate will affect crop production and may jeopardise food security. One important consequence of climate change in this respect is that it is likely to alter interactions between plants and their pathogens (Chaloner et al., 2021; Qiu et al., 2022; Raza and Bebber, 2022; Singh et al., 2023). A recent review of 45 studies on model crop species found that in 55% of cases, disease resistance was negatively affected by elevated temperature (Desaint et al., 2021). This may be driven by changes within the plant, e.g. warmer temperatures may compromise plant basal resistance (Qiu et al., 2022), or by changes within the pathogen, e.g. elevated temperatures can increase pathogen growth rates and reproduction (Vaumourin and Laine, 2018). Pathogen-plant interactions can also be altered due to shared molecular mechanisms underlying virulence and heat tolerance. For example, in rice blast (*Pyricularia oryzae*), genes involved in virulence also influence heat tolerance, (Li et al., 2011). In addition, temperature can alter sensitivity of plant pathogens to fungicides (Lurwanu et al., 2021). Predicting how these interactions will change requires a detailed understanding of how plants and their pathogens respond to changes in temperature as well as knowledge of the molecular mechanisms underlying their response to temperature stress. Increased knowledge of the genes involved in a pathogens’ response to temperature stress may also provide insights into the vulnerabilities of pathogens and potentially identify novel targets for control.

Several molecular responses to temperature stress, particularly higher temperatures, have been well characterized in fungi (e.g. (Bakar et al., 2020; Dunayevich et al., 2018; Richter et al., 2010). In fungi, as in all Eukaryotes, small increases in temperature can cause changes in the three dimensional structure of enzymes and proteins, including unfolding, entanglement, and non-specific aggregation of proteins, and can also affect RNA splicing, DNA replication, cellular membranes and the cytoskeleton (Richter et al., 2010). Repairs to the functional structures of proteins to maintain cell homeostasis are achieved by interactions with molecular chaperones, often referred to as heat shock proteins (HSPs), that prevent aggregation of misfolded proteins and promote efficient protein folding. Although HSPs were first discovered in studies of response to heat shock, these molecular chaperones play key roles in the general stress response, and the initiation of the heat shock response is not necessarily due to changes in temperature, but rather due to the detection of unfolded proteins (Richter et al., 2010; Zheng et al., 2016).

Temperature stress response pathways can overlap with pathways associated with osmotic and oxidative stress. Heat stress can increase damage from oxidative stress caused by the accumulation of reactive oxygen species (ROS), and induce osmotic stress caused by osmotic pressure changes (Xiao et al., 2022). The mitogen-activated protein kinase (MAPK) pathways are important in mediating responses to temperature, oxidative and osmotic stress in fungi. In yeast, there are five MAPK pathways, two of which are activated and interact under temperature stress, including; i) the high-osmolarity glycerol (HOG) pathway (Dunayevich et al., 2018; Winkler et al., 2002) and; ii) the cell wall integrity (CWI) pathway (Sanz et al., 2017). In yeast, heat stress triggers the release of glycerol by activation of CWI; this causes a loss of turgor pressure which stimulates the HOG1 pathway (Dunayevich et al., 2018). In *Zymoseptoria tritici, Pbs2,* a HOG1 pathway gene, was identified as a possible candidate gene for thermal sensitivity (Lendenmann et al., 2016) and heat stress has been shown to trigger intracellular osmotic stress, which triggers the CWI and the HOG pathway and results in hyphal growth (Francisco et al., 2023). These studies demonstrate that both molecular chaperones and MAPK kinases are integral to fungal response to temperature.

Here we use quantitative trait loci (QTL) mapping in *Zymoseptoria tritici* to identify genomic regions associated with responses to high and low temperature stress, enabling a search for candidate genes associated with thermal adaptation. In *Z. tritici*, QTL mapping, genome-wide association studies (GWAS) and transcriptomic studies have already made important contributions to our understanding of the molecular mechanisms underlying fungal response and adaptation to temperature. One of the first QTL studies in *Z. tritici* investigated how colony growth is affected by temperatures that are cooler (15°C) or warmer (22°C) than the optimal growth temperature. This study identified a cool temperature sensitivity QTL on chromosome 10 that contained the candidate gene *Pbs2*, a MAPKK involved in the HOG pathway (Lendenmann et al., 2016). Two GWAS studies analyzed SNP associations using phenotypic data acquired at different temperatures. The first GWAS used 145 global strains to identify many significant SNPs associated with colony size and melanisation at 15°C and 22°C, hereafter referred to as the trait-based GWAS or “traitGWAS” (Dutta et al., 2023). The second used a larger global panel of 411 strains and measured growth traits in liquid media across a broader temperature range (12°C, 15°C, 18°C, 22°C and 25°C) to identify SNPs and candidate genes associated with thermal adaptation, hereafter called “adaptGWAS” (Miñana-Posada et al., 2024). A third GWAS study used a genotype-by-environment association approach (hereafter called “geaGWAS”), based on bioclimatic data obtained from the sampling locations of over 1000 global strains (Feurtey et al., 2023). This study found many significant SNP associations and highlighted promising candidate genes for temperature adaptation.

### Aims

The aim of this study was to identify large effect loci associated with responses to high and low temperature in *Z. tritici* and identify possible candidate genes within the identified QTL intervals. To find the most promising candidate genes to include in future functional validation experiments, we first focussed on the QTL intervals that were unique to temperature stress environments, excluding QTL associated with the same traits measured in the benign temperature environment (18°C). We then used information about gene function, paying attention to known temperature-response genes (Temp-genes, Table S1), GO enrichment, gene variation between the parents, and genes identified in the earlier GWAS analyses to identify the most promising candidate genes.

## Results

### Effect of temperature on growth and melanisation

A total of 209,258 and 194,536 colonies were phenotyped from the 1A5×1E4 and 3D7×3D1 crosses, respectively (see Table S2 for details). Under both hot and cold stress, colony radius was reduced compared to the benign temperature (18°C) in both crosses (Figure 1, Table S2). The colony melanisation was also lower under temperature stress, except at 8 dpi in 27°C in the 3D7×3D1 cross. We can only compare growth and melanisation rate from 8-12dpi between 27°C and 18°C in the 3D7×3D1 cross. We found that growth rate and melanisation rate were both significantly lower under heat stress (Table S2). All pairwise correlations (Pearson, non-parametric) between independent measurements are provided in Tables S3 and S4. We identified many correlations between independent measurements (Table S3 and S4), after excluding traits that are statistically or functionally coupled, including: correlations between two time points of the same trait (i.e. radius at 8 dpi and 12 dpi in the control), correlations between a rate and the corresponding time point measurements (i.e. grey value at 12 dpi and melanisation rate in the control), or correlations between rates and tolerance measurements.

**Figure 1.**
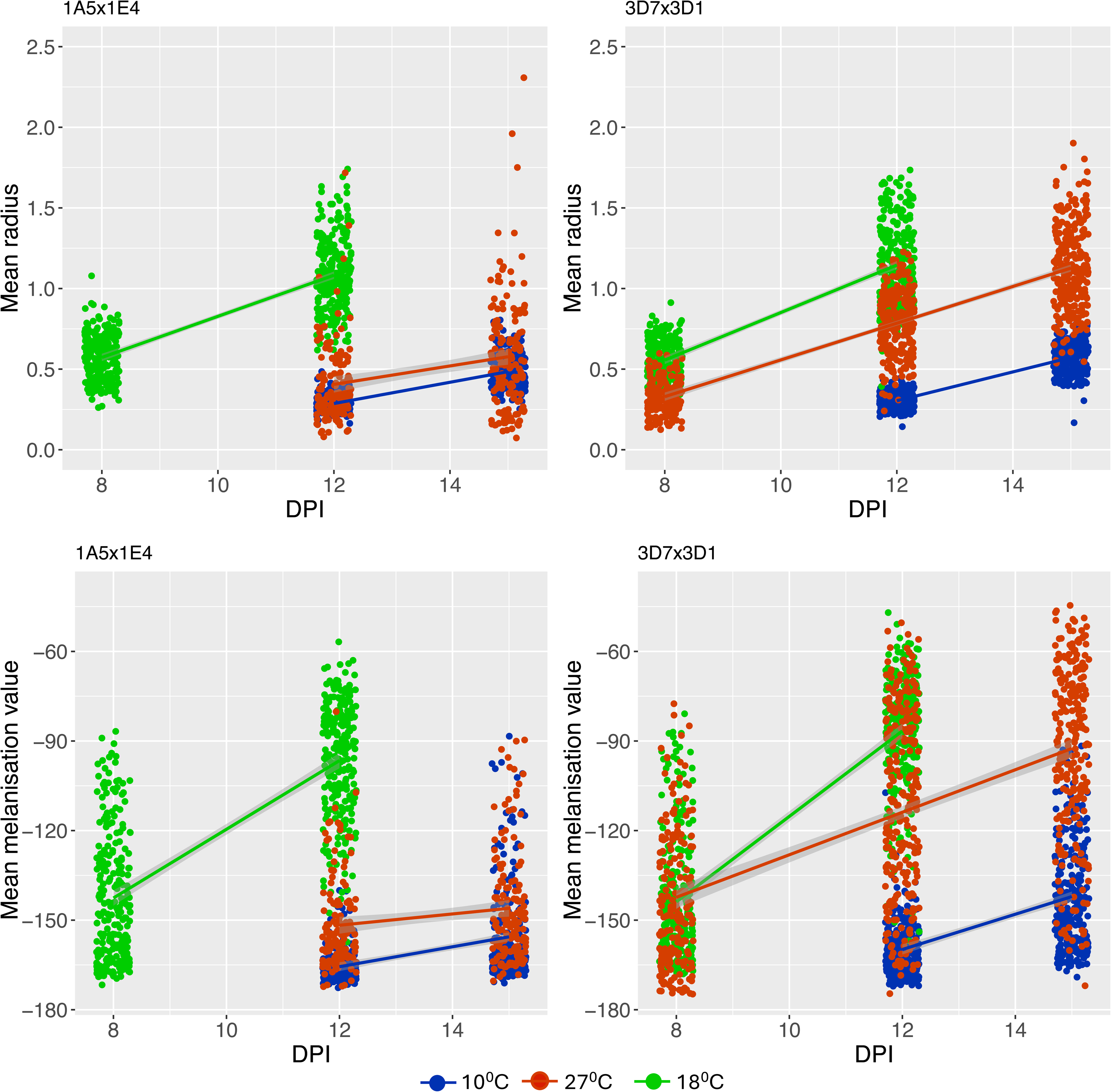
Colony radius and melanisation at three temperatures plotted against days post inoculation (DPI). Colonies grown at high (27°C, red points) or low (10°C, blue points) temperatures had a smaller colony radius than those grown at the 18°C, and were less melanized (panels c, d). Melanization value – inverse of the grey value.

### QTL results

Parent and progeny isolates used in this study are from two crosses between Swiss strains and SNP genotype data for each set of progeny were obtained from a RAD sequence dataset previously produced in our lab (first described in (Lendenmann et al., 2014). Linkage maps built using these SNPs were described previously (Stapley and McDonald, 2023). After filtering and map building, 63181 SNPs were available for analysing the 3D7×3D1 cross and 32806 SNPs were available for the 1A5×1E4 cross (Table S5). We found several QTL associated with growth and melanisation traits (Table 1, Figure 2 and 3). In the 3D7×3D1 cross, significant QTL were found for all but one trait (Growth rate heat tolerance 8-12dpi), and in the 1A5×1E4 cross we found significant QTL for all but four traits (Grey value in 27℃ at 12 dpi, Grey value in 27℃ at 15 dpi, Growth rate 10℃ 12-15 dpi, Melanisation rate 27℃ 12-15 dpi). We separated QTL into those that were uniquely associated with temperature stress from those that were shared across benign and temperature stress environments as described in the methods. In the 3D7×3D1 cross, eight unique temperature QTL (3D7.TQTL) intervals were found (Table 2, Figure 2), and in the 1A5xE4 cross three intervals (1A5.TQTL) were found (Table 2, Figure 3). For full details of these intervals see Table S6 and S7. Within these intervals, interesting genes were identified by combining information from multiple sources, including gene function, sequence variation between parents, GO enrichment, RNAseq data and the presence of SNPs identified in previous GWAS analyses (Dutta et al., 2023; Feurtey et al., 2023).

**Figure 2.**
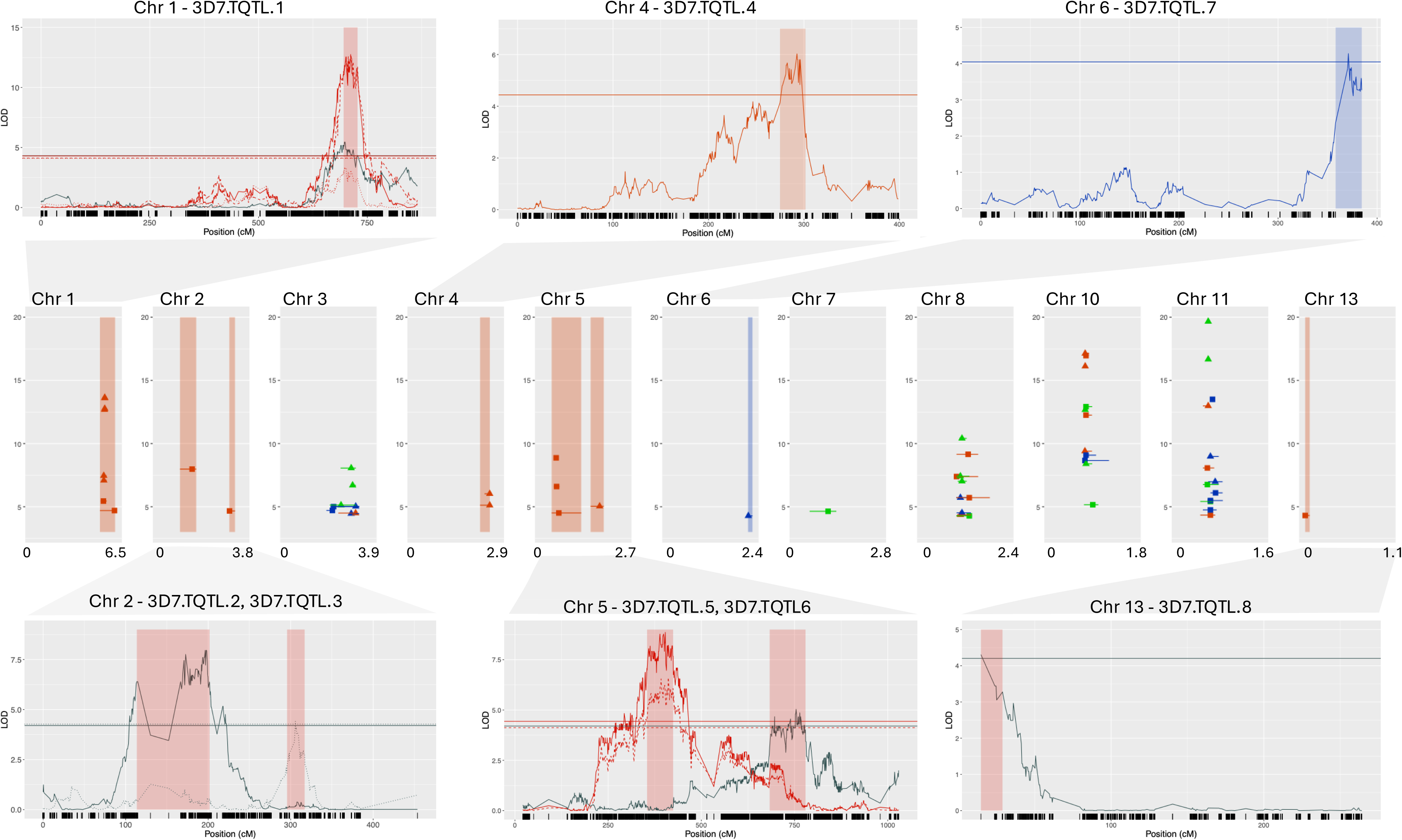
Centre plot shows the genomic location of all significant QTL intervals and the peak LOD value for that interval for the 3D7×3D1 cross. The y-axis is LOD and y-axis genomic position in Megabases (Mb). The shading indicates the unique temperature QTL intervals (3D7.TQTL), red for 27°C, blue for 10°C. Colours indicate if the QTL were associated with traits measured in control (green, 18°C), low temperature (blue, 10°C) or high (red, 27°C) temperature environments, and symbols indicate if the trait is a size (circle) or melanin (triangle) related trait. For each TQTL the detailed LOD profile is provided above or below the centre plot. Some TQTL have multiple traits mapping to the overlapping interval. For plotting we only included one time point trait (solid line), one tolerance trait (dashed line) and one rate trait (dotted). Line colour relates to trait, red: size-related traits at 27°C, blue: size-related traits at 10°C, black: melanin-related trait in any temperature. Shading shows the 95% confidence interval of the QTL, the shading colour relates to the temperature: red for 27°C, blue for 10°C. The x-axis is LOD and y-axis chromosome position in centimorgans (cM), dark lines on the x-axis represent SNP positions.

**Figure 3.**
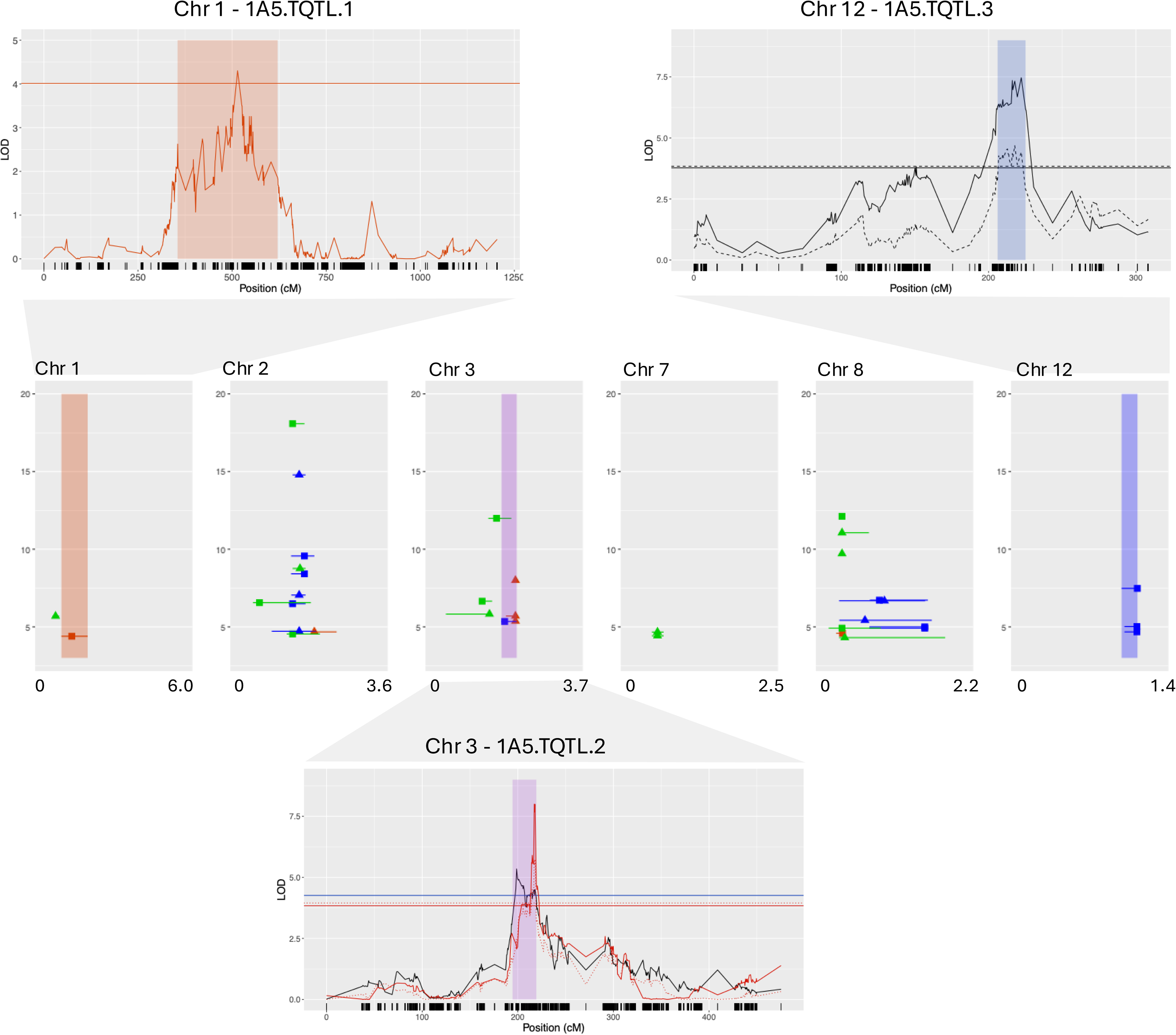
Centre plot shows the genomic location of all significant QTL intervals and the peak LOD value for that interval for the 1A5×1E4 cross. The y-axis is LOD and y-axis genomic position in Megabases (Mb). The shading indicates the unique temperature QTL intervals (1A5.TQTL), red for 27°C, blue for 10°C, purple for both. Colours indicate if the QTL were associated with traits measured in control (green, 18°C), low temperature (blue, 10°C) or high (red, 27°C) temperature environments, and symbols indicate if the trait is a size (circle) or melanin (triangle) related trait. For each TQTL the detailed LOD profile is provided above or below the centre plot. Some TQTL have multiple traits mapping to the overlapping interval. For plotting we only included one time point trait (solid line), one tolerance trait (dashed line) and one rate trait (dotted). Line colour relates to trait, red: size-related traits at 27°C, blue: size-related traits at 10°C, black: melanin-related trait in any temperature. Shading shows the 95% confidence interval of the QTL, the shading colour relates to the temperature: red for 27°C, blue for 10°C, purple for both. The x-axis is LOD and y-axis chromosome position in centimorgans (cM), dark lines on the x-axis represent SNP positions.

**Table 1.** Summary of the QTL mapping results for all traits. Details for each trait are as follows: 1) data type indicates if the measurement was made at a single time point (days post inoculation, DPI) or if it was calculated across two time for details); 2) temperature indicates the temperature at which the colonies were grown, including control (180C), cold stress (100C) and heat stress (270C), or were calculated as a ratio (tolerance) between the cold or heat stress and the growth at 100C vs.180C); 3) measurement type was colony radius or grey value (an indicator of melanization) or the change in radius or melanization over time (growth rate or melanisation rate). For example, in row 1 colony radius was m with the trait measured. QTL columns indicate the chromosomes on which significant QTL peaks were found for each trait in the 3D7×3D1 and 1A5×1E4 crosses. The h^2^ columns show the estimated heritability (h2) for each trait in each c

| Trait full name | No. Isolates<br>3D7x3D1 | QTL 3D7x3D1 | h2 | No. Isolates<br>1A5x1E4 | QTL 1A5x1E4 | h2 |
| --- | --- | --- | --- | --- | --- | --- |
| Colony radius in 10°C at 12 dpi | 263 | 6 | 0.30 | 243 | 2,8 | 0.55 |
| Colony radius in 10°C at 15 dpi | 264 | 8,10 | 0.39 | 225 | 2,8 | 0.43 |
| Colony radius in 18°C at 8 dpi | 261 | 3,8,10 | 0.53 | 251 | 2,3,8 | 0.56 |
| Colony radius in 18°C at 12 dpi | 261 | 3,7,8,10,11 | 0.62 | 250 | 7,8 | 0.48 |
| Colony radius in 27°C at 8 dpi | 202 | 1,4 | 0.58 | Not Measured | Not Measured | NA |
| Colony radius in 27°C at 12 dpi | 258 | 1,4 | 0.42 | 141 | 3 | 0.33 |
| Colony radius in 27°C at 15 dpi | 258 | 1,3 | 0.36 | 149 | 2 | 0.43 |
| Grey value in 10°C at 12 dpi | 263 | 8,10 | 0.55 | 244 | 2,8,12 | 0.48 |
| Grey value in 10°C at 15 dpi | 264 | 8,10,11 | 0.51 | 225 | 2,8,12 | 0.47 |
| Grey value in 18°C at 8 dpi | 261 | 8,10,11 | 0.42 | 251 | 1,2,3,8 | 0.63 |
| Grey value in 18°C at 12 dpi | 261 | 11 | 0.22 | 250 | 2,8 | 0.44 |
| Grey value in 27°C at 8 dpi | 202 | 11 | 0.81 | Not Measured | Not Measured | NA |
| Grey value in 27°C at 12 dpi | 258 | 1,11 | 0.64 | 139 | No QTL | 0.40 |
| Grey value in 27°C at 15 dpi | 258 | 1,3,11 | 0.56 | 149 | No QTL | 0.22 |
| Growth rate 10°C 12-15 dpi | 263 | 10 | 0.40 | 218 | No QTL | 0.28 |
| Growth rate 18°C 8-12 dpi | 258 | 3,8,11 | 0.61 | 249 | 7,8 | 0.33 |
| Growth rate 27°C 8-12 dpi | 199 | 1,3 | 0.33 | Not Measured | Not Measured | NA |
| Growth rate 27°C 12-15 dpi | 256 | 1 | 0.21 | 127 | 2 | 0.30 |
| Melanisation rate 10°C 12-15 dpi | 263 | 8,10 | 0.35 | 219 | 2,8,12 | 0.47 |
| Melanisation rate 18°C 8-12 dpi | 258 | 10 | 0.28 | 249 | 2,3 | 0.54 |
| Melanisation rate 27°C 8-12 dpi | 199 | 11 | 0.26 | Not Measured | Not Measured | NA |
| Melanisation rate 27°C 12-15 dpi | 256 | 3,11 | 0.31 | 125 | No QTL | 0.28 |
| Colony radius cold tolerance at 12 dpi | 259 | 3,11 | 0.44 | 243 | 2 | 0.33 |
| Colony radius heat tolerance at 8 dpi | 198 | 1,5,10 | 0.60 | Not Measured | Not Measured | NA |
| Colony radius heat tolerance at 12 dpi | 254 | 1,5,8,10,11 | 0.57 | 141 | 3 | 0.35 |
| Grey value cold tolerance at 12 dpi | 259 | 11 | 0.22 | 244 | 3 | 0.40 |
| Grey value heat tolerance at 8 dpi | 198 | 11 | 0.43 | Not Measured | Not Measured | NA |
| Grey value heat tolerance at 12 dpi | 254 | 11 | 0.36 | 139 | 8 | 0.39 |
| Growth rate heat tolerance at 8-12 dpi | 194 | No QTL | 0.48 | Not Measured | Not Measured | NA |
| Melanisation rate heat tolerance at 8-12 dpi | 194 | 8,11 | 0.26 | Not Measured | Not Measured | NA |
| Survival (colony presence/absence) in 27°C at 12 dpi | Not Measured | Not Measured | NA | 248 | 1 | 0.48 |

**Table 2.** Summary of the unique temperature QTL intervals, including the TQTL ID, associated chromosome, temperature stress (heat/cold/both) and the number of traits associated with this locus (M=melanization, S=colony size, B=both includes: start and stop wpositions and interval size in Kilobases, maximum LOD, number of genes, number of Temp-genes (Tgenes), Temp-gene names, the p-value indicating if the interval contained more heat shock proteins (HSPs) than annotations, the number of genes overlapping with the traitGWAS SNPs, the geaGWAS SNPs or the adaptGWAS.

| QTL.ID | Chr | Temp | No.traits (type) | Start | Stop | Interval | Max LOD | No. genes | No.<br>Tgenes | Temp-genes | HSP enrich | GO enrich | geaGWAS | traitGWAS | adaptGWAS |
| --- | --- | --- | --- | --- | --- | --- | --- | --- | --- | --- | --- | --- | --- | --- | --- |
| 3D7.TQTL.1 | 1 | Heat | 8 (B) | 5189.86 | 6311.30 | 1121.44 | 13.15 | 391 | 5 | 2xHSP, 1xMAPKK Ssk2/Ssk22, 2xC2H2 | p=0.27 | 19 | 47 | 1 (RCA) | 5 (D2.12C, eAUC12C) |
| 3D7.TQTL.2 | 2 | Heat | 1 (M) | 1004.51 | 1626.25 | 621.74 | 8.05 | 250 | 7 | 2xC2H2, 3xHSP, 2xPTP | p=0.32 | 15 | 8 | 0 | 1 (x75.12C) |
| 3D7.TQTL.3 | 2 | Heat | 1 (M) | 3169.15 | 3428.44 | 259.30 | 4.39 | 67 | 1 | 1xcAMP | NA | 5 | 2 | 0 | 0 |
| 3D7.TQTL.4 | 4 | Heat | 3 (S) | 2455.95 | 2646.71 | 190.77 | 6.12 | 124 | 1 | 1xHSP | p=0.33 | 3 | 18 | 1 (M15C) | 0 |
| 3D7.TQTL.5 | 5 | Heat | 3 (S) | 399.93 | 1237.36 | 837.43 | 8.95 | 332 | 3 | 2xHSP, 1xPTP | p=0.04 | 13 | 37 | 0 | 0 |
| 3D7.TQTL.6 | 5 | Heat | 1 (M) | 1661.50 | 2062.60 | 401.10 | 5.03 | 21 | 6 | 4xHSP, 3xC2H2 | p=0.06 | 8 | 3 | 0 | 0 |
| 3D7.TQTL.7 | 6 | Cold | 1 (S) | 2260.91 | 2383.12 | 122.21 | 4.29 | 40 | 0 |  | NA | 4 | 1 | 0 | 0 |
| 3D7.TQTL.8 | 13 | Cold | 1 (M) | 23.95 | 82.58 | 58.62 | 4.39 | 124 | 0 |  | NA | 1 | 1 | 0 | 0 |
| 1A5.TQTL.1 | 1 | Heat | 1 (S) | 1037.31 | 1949.98 | 912.67 | 4.39 | 439 | 5 | 1xOpy2, 4xHSP, 1xMAP, 1xPTP, 1xcAMP(PKA) | p=0.08 | 19 | 23 | 0 | 0 |
| 1A5.TQTL.2 | 3 | Both | 4 (B) | 1843.71 | 2251.85 | 408.13 | 8.00 | 114 | 2 | 1xSho1, 1xPTP | NA | 3 | 7 | 1 (RCA) | 0 |
| 1A5.TQTL.3 | 12 | Cold | 3 (M) | 1070.06 | 1234.96 | 164.90 | 7.48 | 61 | 2 | 1xC2H2, 1xPTP | NA | 2 | 3 | 0 | 0 |

KEY: RCA: Relative Colony Area 14-22C, M15C: Melanin 15C, D2.12C: day2\_12C, x75.12C: xq75total\_12C

### GO annotations and HSP enrichment in TQTL

We grouped all genes from TQTL intervals according to the stress (heat/cold) and the trait (colony size/melanisation) to create gene sets. In both crosses we had three gene sets for analysis and the results are provided in Tables S8 and S9. The largest enriched GO annotation for 3D7×3D1, in terms of the highest number of significant genes, was “zinc ion binding” (GO:0008270). The interval 3D7.TQTL.1 had 12 genes with this GO annotation, and the interval 3D7.TQTL.2 had 13 genes with this annotation. In the 1A5×1E4 cross, the two GO annotations with the highest number of significant terms were “protein phosphorylation” (GO:0006468) and “protein kinase activity” (GO:0004672). Fourteen genes had these GO annotations in the interval 1A5.TQTL.1. One interval, 3D7.TQTL.5, as significantly enriched for QTLs (Table 2), having more than the number of HSPs observed in 10,000 random QTL intervals on the same Chromosome (p=0.04). Two other TQTL intervals had four HSPs, one in each cross (3D7.TQTL.6 and 1A5.TQTL.1). The probability of observing four HSPs in an interval on each of those two respective chromosomes was 0.06 and 0.08 (Table 2), thus although it was uncommon (p<0.10), it was not significant at p<0.05.

### Most promising candidate genes associated with temperature stress in 3D7×3D1

The 3D7.TQTL interval with the highest LOD was on chromosome 1 (3D7.TQTL.1) and it was associated with six colony size-related traits (colony radius in 27°C at all time points, colony radius tolerance at 8 and 12dpi and growth rate between 8-12dpi) and two melanisation-related traits (mean grey value at 27°C at 12 and 15dpi). The size-related QTL had a higher Lod (>10) than the melanin traits (Figure 2). The largest significant interval spanned 1121 Kb and contained 391 genes. Of these 391 genes, 46 contained geaGWAS-SNPs, one traitGWAS-SNP, and 19 had enriched GO terms (Table 2, Table S10). We provide a summary of several genes of interest (Table 3) and highlight next the four most promising candidate genes in this interval.

**Table 3.** Details connected with the most promising candidate genes found within TQTL intervals. This includes: cross, TQTL ID, chromosome (Chr), gene ID, protein length (Prot.ln), the bioclimatic variable associated with any GWAS-SNP association with SNP in thermal adaption GWAS (adapGWAS), if the gene had an enriched GO annotation,, (Protein Sim.) between the parents, if the gene was differentially expressed *in vitro* or *in vivo*, IPO323 gene ID, IPO323 protein le protein family domain within the gene.

| TQTL ID | Chr | ZT3D7 Gene ID | ZT3D7 Prot.In | geaGWAS | traitGWAS | adaptGWAS | GO annotation enriched | ZT3D1 Gene ID | ZT3D1 Prot.In | Protein sim (3D7, 3D1) | High Impact Var. | DEG invitro | DEG invivo | IPO323.geneID | IPO323 Prot.In | Protein sim (IPO, 3D7) | Protein sim (IPO, 3D1) | Predicted domain |
| --- | --- | --- | --- | --- | --- | --- | --- | --- | --- | --- | --- | --- | --- | --- | --- | --- | --- | --- |
| 3D7.TQTL.1 | 1 | ZT3D7_G1860 | 1654 | NA | NA | NA | No | ZT3D1_G1822 | 1618 | 91.88 | 3 | Yes | Yes | ZtiIPO323_019890 | 1688 | 94.92 | 93.06 | Histidine kinase/HSP90-like ATPase |
| 3D7.TQTL.1 | 1 | ZT3D7_G1950 | 810 | BIO6 | NA | NA | Yes | ZT3D1_G1910 | 867 | 92.96 | 0 | Yes | Yes | ZtiIPO323_020880 | 300 | 100 | 87 | probable general stress response pr |
|  |  |  |  |  |  |  |  |  |  |  |  |  |  | ZtiIPO323_020890 | 360 | 99 | 99 | Alcohol dehydrogenase GroES-like c<br>alcohol dehydrogenase |
| 3D7.TQTL.1 | 1 | ZT3D7_G1951 | 1385 | NA | NA | NA | No | ZT3D1_G1911 | 1419 | 99.93 | 0 | Not sig. | NA | ZtiIPO323_020900 | 1419 | 99.86 | 99.93 | Protein kinase domain, Mitogen-act |
| 3D7.TQTL.1 | 1 | ZT3D7_G2013 | 581 | NA | RCA | NA | No | ZT3D1_G1975 |  | 88.66 | 1 | Yes | Not sig. | ZtiIPO323_021600 | 581 |  |  | No predicted function |
| 3D7.TQTL.1 | 1 | ZT3D7_G2052 | 520 | NA | NA | eAUC12C | No | ZT3D1_G2016 | 520 | 99.23 | 0 | Not sig. | Not sig. | ZtiIPO323_022050 | 520 | 99.42 | 98.65 | Ammonium Transporter Family |
| 3D7.TQTL.4 | 4 | ZT3D7_G5373 | 449 | NA | M15C | NA | No | ZT3D1_G5304 | 449 | 99.78 | 0 | Not sig. | Yes | ZtiIPO323_058350 | 362 | 100 | 99.72 | MOSC N-terminal beta barrel domai |

| TQTL ID | Chr | ZT1A5 Gene ID | ZT1A5 Prot.In | geaGWAS SNP | traitGWAS SNP | adaptGWAS SNP | GO annotation enriched | ZT1E4 Gene ID | ZT1E4 Prot.In | Protein sim (1A5, 1E4) | High Impact Var. | DEG invitro | DEG invivo | IPO323.geneID | IPO323 Prot.In | Protein sim (IPO, 1A5) | Protein sim (IPO, 1A4) | Predicted domain |
| --- | --- | --- | --- | --- | --- | --- | --- | --- | --- | --- | --- | --- | --- | --- | --- | --- | --- | --- |
| 1A5.TQTL.1 | 1 | ZT1A5_G381 | 1547 | NA | NA | NA | No | ZT1E4_G375 | 1547 | 99.55 | 1 | Not sig. | Not sig. | ZtiIPO323_004060 | 1513 | 97.29 | 97.48 | Histidine kinase/HSP90-like ATPase |
| 1A5.TQTL.1 | 1 | ZT1A5_G420 | 414 | NA | NA | NA | Yes | ZT1E4_G414 | 414 | 100 | 0 | Not sig. | Not sig. | ZtiIPO323_004450 | 414 | 100 | 100 | Protein kinase domain, Mitogen-act |
| 1A5.TQTL.1 | 1 | ZT1A5_G473 | 185 | NA | NA | NA | No | ZT1E4_G467 | 185 | 96.22 | 0 | Not sig. | Not sig. | ZtiIPO323_005050 | 187 | 95.14 | 96.24 | PTP, Low molecular weight phospho |
| 1A5.TQTL.1 | 1 | ZT1A5_G491 | 442 | BIO5 | NA | NA | No | ZT1E4_G485 | 453 | 90.27 | 1 | Not sig. | Not sig. | ZtiIPO323_005250 | 459 | 91.94 | 88.06 | WD40-repeat-containing domain su |
| 1A5.TQTL.1 | 1 | ZT1A5_G492 | 476 | BIO5, BIO10 | NA | NA | No | ZT1E4_G486 | 476 | 99.79 | 0 | Not sig. | Not sig. | ZtiIPO323_005260 | 476 | 99.58 | 99.79 | Opy2 protein |
| 1A5.TQTL.1 | 1 | ZT1A5_G634 | 1178 | BIO3 | NA | NA | No | ZT1A5_G632 | 1176 | 97.28 | 0 | Not sig. | Not sig. | ZtiIPO323_006800 | 1176 | 97.87 | 97.88 | Histidine kinase/HSP90-like ATPase |
| 1A5.TQTL.2 | 3 | ZT1A5_G4068 | 329 | NA | NA | NA | No | ZT1E4_G4135 | 329 | 100 | 0 | Not sig. | Not sig. | ZtiIPO323_044480 | 329 | 100 | 100 | SH3 domain, Sho1, SH3 domain, Hig |
| 1A5.TQTL.2 | 3 | ZT1A5_G4089 | 549 | NA | RCA | NA | No | ZT1E4_G4154 | 549 | 100 | 0 | Yes | Not sig. | ZtiIPO323_044730 | 549 | 100 | 100 | Cytochrome P450 |
| 1A5.TQTL.3 | 12 | ZT1A5_G11001 | 779 | NA | NA | NA | No | ZT1E4_G11018 | 779 | 99.49 | 0 | Yes | Not sig. | ZtiIPO323_120430 | 780 | 99.48 | 99.74 | Zinc finger, C2H2 type |
| 1A5.TQTL.3 | 12 | ZT1A5_G11023 | 355 | NA | NA | NA | No | ZT1E4_G11044 | 355 | 100 | 0 | Yes | Yes | ZtiIPO323_120710 | 390 | 100 | 100 | Tyrosine phosphatase family (PTP) |

The first gene encodes a HSP90-like ATPase superfamily domain (ZT3D7_G1860) that we consider a promising candidate because it varied between the two parents, including three high impact variants (two “frameshift variants”, one “stop gained”) likely to affect protein function. Under benign temperature the gene is differentially expressed *in vitro* and *in vivo* (Table 3). This gene was in the high Lod peak associated with size-related traits.

The second promising candidate gene is a “probable general stress response protein *Whi2”*. This gene is not annotated in the 3D7 and 3D1 genomes, but it is annotated in the most recent IPO323 annotation (gene ID: ZtIPO323_020880) and is orthologous with part of ZT3D7_G1950, which has an enriched GO term (zinc ion binding, GO:0008270) and a geaGWAS-SNP for Minimum Temperature of Coldest Month (BIO6) (Table 3). The gene ZtIPO323_020880, predicted to encode *Whi2* has 100% protein conservation with part of ZT3D7_G1950, but only 87% protein conservation with part of ZT3D1_G1910 (Table 3).

Based on the 3D7 genome annotation this gene contains an alcohol dehydrogenase GroES-like domain (GroES is a HSP for *E. coli*) and a zinc-binding dehydrogenase domain. After blasting the alcohol dehydrogenase protein (ZtIPO323_020890) against the 3D7 and 3D1 genomes, we found only one mismatch in ZT3D1_G1910 and two in ZT3D7_G1950 among the 360 amino acids encoded by this gene. The genomic region containing ZT3D7_G1950 and ZT3D1_G1910 needs to be reannotated in these respective reference genomes.

The third (ZT3D7_G1951), is immediately adjacent to ZT3D7_G1950 and ZT3D1_G1910 and is a MAPKKK *SSk2/Ssk22* (*ZtSsk2*) gene, which is part of the HOG1 pathway (Dunayevich et al., 2018). *ZtSsk2* is homologous across the three relevant genomes (ZT3D7_G1951, ZT3D1_G1911, ZtIPO323_020900) with little protein variation, only one amino acid difference between ZT3D1_G1911 and ZT3D7_G1951 (Alanine-Valine), and there no significant differences in gene expression under benign temperatures (Table 3).

The fourth promising candidate gene in this interval is ZT3D7_G2052, which encodes an ammonium transport family domain. This gene has multiple adaptGWAS SNPs and the protein varies between the parents, with no high impact SNPs, but four SNPs predicted to have moderate effects (https://github.com/jessstapley/QTL-mapping-Z.-tritici/). The gene was upregulated in 3D7 in vitro growing on minimal media.

### Most promising candidate genes associated with temperature stress in 1A5×1E4

On chromosome 1, the 1A5.TQTL.1 was associated with the binary trait Colony growth at 27°C. This interval was large and contained 439 genes (Table 2, Table S11), including 19 genes with enriched GO annotations and 22 genes with a geaGWAS SNP. This interval contains several promising candidates (Table 3) including the HOG1 pathway gene highlighted in the geaGWAS, *Opy2* (ZT1A5_G492). The *Opy2* geaGWAS SNP was associated with Maximum Temperature of Warmest Month (BIO5, -log(p-value) = 7.65) and Mean Temperature of Warmest Quarter (BIO10, -log(p-value) = 7.60). There is one amino acid difference between the parents (Arginine-Serine). The second promising candidate is a HSP gene with a geaGWAS SNP associated with Isothermality (BIO3: mean diurnal temperature range/temperature annual range) (ZT1A5_G634). This protein which is 1176 amino acids long, has 32 amino acids differences between parents and 37 moderate effect variants (missense variants). The second unique temperature QTL 1A5.TQTL.2 for this cross was on chromosome 3. It was associated with both hot and cold stress and contained the HOG1 signalling protein, *Sho1*, (ZT1A5_G4068) (Table 2, 3) as well as a cytochrome P450 gene with a traitGWAS SNP (ZT1A5_G4089). The protein products of these genes were conserved between the parents, and also with the IPO323v2 annotation. No significant differences in the gene expression between parents *in vitro* or *in vivo* at the Sho1 gene, however the cytochrome p450 gene was upregulated *in vitro* (Table S11)

On chromosome 12, there was a cold-stress-QTL interval (1A5.TQTL.3) associated with three melanisation traits (Melanisation rate 10°C 12-15 dpi, Grey value in 10°C at 15 dpi, Grey value in 10°C at 12 dpi). This interval contained 62 genes and overlapped, by a single gene, with a QTL previously identified for the morphological transition from yeast-like growth to hyphal growth at high temperature (Francisco et al., 2023) (Table S11). Within the interval there were two promising candidate genes (Table 3), including ZT1A5_G11001 which has a Zinc finger C2H2 type domain, and ZT1A5_G11023, which has a Protein-tyrosine phosphatase-like (PTP) domain. Both genes showed differential expression between the parents (Table 3). The zinc finger C2H2 type gene produced a different protein in the parents and the IPO323v2 orthologue is predicted to encode the transcriptional regulator CRZ1. CRZ1 was shown to be involved in cell wall integrity and melanisation in *Alternaria alternata* (Yang et al., 2022) and heat-shock response in pathogenic fungi (Roy et al., 2021).

## Discussion

We identified 13 and 11 distinct QTL intervals associated with growth and melanisation in the 3D7×3D1 and 1A5×1E4 crosses, respectively (Figure 2 and 3). Among these, we identified 11 QTL intervals uniquely associated with temperature stress (TQTL), with eight such intervals in the 3D7×3D1 cross and three in the 1A5×1E4 cross (Figures 2 and 3, Table 2). Based on information gathered from multiple sources including previous GWAS analyses and functional annotations, we highlight several promising candidate genes for temperature responses including: HSP90 genes, a *Whi2* gene, a zinc-dependent alcohol dehydrogenase, an ammonium transporter, and two HOG pathway genes *SSk2* and *Opy2*. HSPs were common in the TQTL intervals. There was some evidence of enrichment (Table 2), 3D7.TQTL.5 had significantly (p=0.04) more HSPs than the number of HSPs observed in 10,000 random QTL intervals. Two intervals had 4 HSPs, and although this was not significantly (p<0.05) greater than the number of HSPs observed in 10,000 random QTL intervals, it was uncommon, occurring in only six and eight percent of these randomly generated intervals for 3D7.TQTL.6 and 1A5.TQTL.1 respectively. In a previous QTL study using the same isolates grown *in vitro* under cool-temperature (15°C) stress, no enrichment of HSPs was found (Lendenmann et al., 2016), but our new analyses that included a broader range of temperatures suggests that HSPs may contribute to responses to temperature.

### Candidates for temperature stress response identified in the 3D1×3D7 cross

Heat shock proteins form an integral part of the general stress response in many organisms (Richter et al., 2010). We highlighted a HSP90-like ATPase superfamily gene (ZT3D7_G1860) as a promising candidate. HSP90 is the most common form of HSP in yeast and the most common predicted HSPs in our reference genomes (31 in 3D7, 32 in 1A5). HSP90 interacts with hundreds of proteins under both stressful and stress-free environments (Crunden and Diezmann, 2021). In *Candida albicans*, HSP90s regulate the transition between yeast and filamentous forms as well as virulence (Robbins and Cowen, 2023). The 3D7.TQTL.6 interval had four HSPs, two of which were HSP70 genes (ZT3D7_G6242, ZT3D7_G6240). HSP70s are less common in the genome (11 in 3D7 and 12 in 1A5), but are one of the most conserved HSPs. They are involved in protein folding, both de novo folding and refolding of aggregated proteins, and prevent aggregation of denatured proteins under stress (Richter et al., 2010). In *Z. tritici,* expression of four HSP70 genes was upregulated at the transition stage from biotrophy to necrotrophy (7-11 days post inoculation) in planta (Palma-Guerrero et al., 2016). One of these, Mycgr3G72449 (Zt09_model_5_00606), is the orthologue of ZT3D7_G6242 (100% similarity at the protein level) which is in the 3D7.TQTL.6 interval. Another notable feature of the two HSP70 genes in the 3D7.TQTL.6 interval is that they are very close, separated by only a single gene, which is a Zinc finger C2H2-type gene.

We identified most of the candidate genes in the high temperature response QTL on Chromosome 1 (3D7.TQTL.1) (Table 3). This QTL contained 47 genes with SNPs significantly associated with climatic variables (Feurtey et al., 2023), and was associated with growth at cooler temperatures (relative colony area between 15°C and 22°C (Dutta et al., 2023), and growth at 12°C (Miñana-Posada et al., 2024). In the adaptGWAS, one gene contained seven significant SNPs associated with 12°C growth across a panel of 411 isolates (Miñana-Posada et al., 2024). The gene encodes an ammonium transporter-like protein (Table 3). In contrast to the previous QTL and GWAS studies, our QTL was associated with growth and melanisation under high temperature (27°C) but not cold temperature (10°C). If these candidate genes are related to a response to temperature, then our results, combined with the previous studies, suggest that they are affecting a general temperature response rather than a response limited to either high or low temperatures.

The other genes we highlight as promising candidates are two adjacent genes in 3D7.TQTL.1. One has orthology with a probable general stress response protein *Whi2 (*ZT3D7_G1950) and the other gene encodes a MAP kinase kinase kinase *SSk2/Ssk22* domain (ZT3D7_G1951). ZT3D7_G1950 is a potential candidate because it has an enriched GO term (GO:0008270, Zinc ion binding), has a geaGWAS SNP for Minimum Temperature of Coldest Month (Table 3) and was differentially expressed between the parents – the gene was upregulated in 3D1 compared to 3D7 in minimal media and on Drifter at 28 dpi (Table S10). There is no one-one homology of ZT3D7_G1950 with the most recent IPO323 genome and alignment of this region between the two parents (3D7, 3D1) and IPO323 genome suggests the annotation in 3D7 and 3D1 needs updating. Based on DNA and protein similarity with IPO323, there is a gene encoding *Whi2* and a zinc dependent ADH at this position in 3D7 and 3D1. WHI2 (Whiskey2) is a general stress response factor that can influence the expression of stress-associated genes, nutrient sensing and the transition from biotrophic infection to necrotrophic infection in fungi (Kaida et al., 2002; Shuai et al., 2022). If *Whi2* effects a response to temperature in pathogenic fungi has, to our knowledge not been tested, but in the rice false smut (*Ustilaginoidea virens), UvWhi2* knockout had increased sensitivity to oxidative stress and cell wall stress (Shuai et al., 2022), both of which can be induced heat stress. ADHs are important in growth and metabolism under both aerobic and anaerobic conditions and are known to affect virulence in many fungal genera (Gutiérrez-Corona et al., 2023). The roles played by ADHs under temperature stress in pathogenic fungi remains unclear, however ADHs were shown to be involved in cold stress tolerance in plants (Christie et al., 1991) and the oleaginous alga *Nannochloropsis salina* (Lim et al., 2023).

ADHs also play important roles in tolerance to oxidative stress (Gutiérrez-Corona et al., 2023), which can be induced by temperature stress (Xiao et al., 2022). The other potential candidate, *SSk2,* is involved in the HOG1 pathway and was already shown to play a role in heat stress response in yeast (Dunayevich et al., 2018). In *Z. tritici*, loss of function mutations in *Ssk2* led to greater sensitivity to osmotic and oxidative stress, loss of virulence on a susceptible host (cv. Riband) and increased resistance to fungicides (Blyth et al., 2023). It would be worthwhile to test the responses of *Ssk2* mutants to temperature stress.

### Candidates for temperature stress response in the 1A5×1E4 cross

A notable feature of this cross was that the parent 1E4 and approximately half of the offspring in this cross did not grow under heat stress (27°C). We identified a QTL associated with this binary trait (growth/no-growth at 27°C), 1A5.TQTL.1. Within this QTL interval, there were many Temp-genes, many genes with geaGWAS SNPs and one gene with a traitGWAS SNP. A notable gene with a geaGWAS SNP is *Opy2*. *Opy2* has been shown to be involved in responses to stress and affects virulence in fungi. In *S. cerevisiae, Opy2* transmembrane protein is involved in the *Sho1* branch of the HOG1 pathway during osmotic stress (Smith et al., 2010). It was also shown to play a role in the filamentous growth pathway in response to glucose limitation (Karunanithi and Cullen, 2012). In *Candida albicans*, *CaOpy2* does not play a role in adaptation to osmotic stress, but it is required for *Cek1* phosphorylation and plays an essential role in cell wall stress (Herrero De Dios et al., 2013). *Opy2* was shown to affect virulence in *Metarhizium robertsii,* (Guo et al., 2017) and *Magnaporthe oryzae* (Cai et al., 2022). The protein products of genes encoding *Opy2* in our two parents (ZT1A5_G492 and ZT1E4_G486) were very similar, with only a single amino acid difference (arginine-serine), but *Opy2* expression did not differ between the parents on different media or during infection at benign temperatures (Table S11). In a previous study searching for QTL associated with osmotic stress using the same isolates, one osmotic stress QTL on Chromosome 1 was found and the *Opy2* gene was present in that interval (Stapley and McDonald, 2023). The QTL was associated with three traits (Colony radius KCl tolerance at 12dpi, Growth rate KCl tolerance, Colony radius in KCl at 12dp) but had a rather low LOD (4.03-4.80). The largest interval of these three QTL spanned 5583 Kb and contained over 2000 genes. Because of the relatively low LOD and large interval, we did not consider possible candidates in this interval in the previous study, but focussed instead on a smaller interval that had a larger LOD (Stapley and McDonald, 2023). Taking into account the finding that *Opy2* is present in independent QTL for osmotic and temperature stress, we speculate that heat stress may trigger changes in turgor pressure that can be sensed by the osmosensor *Opy2* and trigger the CWI pathway, as was found in yeast (Dunayevich et al., 2018; Smith et al., 2010).

The 1A5.TQTL.3 on chromosome 12 associated with cold stress partially overlapped with a previously mapped QTL for a heat-induced blastospore-to-hyphae/chlamydospore transition (Francisco et al., 2022). The previous study functionally validated two genes that regulate the blastospore-to-hyphae/chlamydospore transition in response to heat stress in *Z. tritici*, including a novel transcription factor named *ZtMsr1* and a protein phosphatase, *ZtYvh1*. The overlapping region between 1A5.TQTL.3 and the previous interval is small and only contains only one gene (ZT1A5_G11033), not the two genes functionally annotated in the previous study. Our interpretation in this case is that the two traits involve different genes even though the QTLs showed a small overlap.

### QTL intervals associated with traits measured in stressful and benign temperature environments

We found a QTL on chromosome 10 that was associated with colony size and melanisation traits measured in both cold stress and benign environments, as well as colony radius heat tolerance. Because there is an overlap between the intervals identified in stressful and benign environments, this is not a unique temperature stress QTL. However it is worth mentioning because it overlaps with a QTL containing a candidate gene (*Pbs2*) for cold sensitivity previously identified in the 3D7×3D1 cross (Lendenmann et al., 2016). *Pbs2* is upstream of HOG1 and is downstream of *SSk2* and *Opy2*. In the previous QTL study colonies were grown at 15°C and 22°C, thus under a milder cold stress compared to our current study. A QTL associated with oxidative stress tolerance was also found on chromosome 10 in a previous study (Zhong et al., 2021), however that QTL does not overlap with the QTL we identified in this study.

Another notable QTL interval is that on chromosome 11. This was previously found to be associated with melanisation and colony size traits in stressful and benign environments (Lendenmann et al., 2014; Stapley and McDonald, 2023) in the 3D7×3D1 cross. Regulation of the *Zmr1* transcription factor in this interval was shown to affect melanin production and is thought to explain most of the phenotypic variance associated with this QTL (Krishnan et al., 2018). We again found that multiple melanisation- and growth-related traits measured under hot and cold stress were associated with this QTL interval (Figure 2, Table S6) and believe that the previous interpretations also apply in these cases.

In the 1A5×1E4 cross, we found a large QTL interval on chromosome 8 that included 11 traits, including colony size and melanisation traits under temperature stress and benign temperature environments (Table S7). The same QTL was found in previous studies mapping QTL for oxidative (Zhong et al., 2021) and osmotic stress (Stapley and McDonald, 2023), and similar to results of this study, the QTL was associated with traits measured in both benign an stressful environment. This genomic regions likely contains genes that affect intrinsic growth of colonies under a wide range of environments and/or genes with pleiotropic effects.

### Notable temperature stress genes identified via GO annotation enrichment

A notable GO annotation was the zinc ion binding (GO:0008270) term. Many genes in unique temperature QTL had this annotation, and it was enriched amongst the gene set associated with melanisation related traits under heat stress. Zinc finger proteins are one of the largest transcription factor families in eukaryotes (Ding et al., 2022). Genes with this annotation encode many proteins, including C2H2-type zinc-finger proteins found in one of our Temp-genes, and are considered master regulators of stress response in plants (ref), mushrooms (Hou et al 2020), and other eukaryotes (). Genes encoding fungal Zn(2)-Cys(6) binuclear cluster proteins, which bind two zinc atoms with a DNA-binding domain consisting of six cysteine residues, also have this GO annotation, which is found over 100 times in our reference genomes (3D7:102, 1A5: 125). In mushrooms these have been shown to be differentially expressed under cold and heat stress (ref). In *Z. tritici*, the Zn(2)-Cys(6) binuclear cluster domain gene ZtMsr1 is involved in the blastospore-to-hyphae/chlamydospore transition that is triggered by heat stress. Deletion of ZtMsr1 in the 1A5 strain activated hyphal growth, whereas the insertion of the 1A5-ZtMsr1 allele into the 1E4 strain (in which ZtMsr1 is naturally disrupted by a TE), induced chlamydospore formation at 27◦C (Francisco et al., 2022). This example illustrates how zinc finger proteins can influence important fungal traits under temperature stress.

## Methods

### QTL mapping crosses

Parent and progeny isolates used in this study are from two crosses between Swiss strains that were described previously (Lendenmann et al., 2014). The original choice of strains to cross was based upon seven measured traits for each strain, including in planta virulence, pycnidia size and density, and in vitro growth rates under different temperatures and in the presence of fungicides (Zhan et al., 2005). The two crosses yielded 700 offspring to use for QTL mapping. The four parents were later phenotyped under other environmental stressors (e.g. reactive oxygen (Zhong et al., 2021)) and at two temperatures (15°C and 22°C) (Lendenmann et al., 2016). The cross between ST99CH1A5 and ST99CH1E4 (herein referred to as 1A5 and 1E4 respectively) produced 341 progeny. The cross between ST99CH3D7 and ST99CH3D1 (herein referred to as 3D7 and 3D1 respectively) produced 359 progeny.

### Genotyping

SNP genotype data for each set of progeny were obtained from a RAD sequence dataset previously produced in our lab (first described in (Lendenmann et al., 2014). In brief, the genome was cut using the restriction enzyme *Pst*l and the libraries were sequenced on an Illumina HiSeq2000 with paired-end sequencing. Complete genome sequences of the parental strains (NCBI Biosample: SRS383146 (ST99CH3D1), SRS383147 (ST99CH3D7), SRS383142 (ST99CH1A5), and SRS383143 (ST99CH1E4)) were used to SNP genotype the parents and offspring (Croll et al., 2013).

RADseq processing, variant discovery and linkage maps are described in detail at https://github.com/jessstapley/QTL-mapping-Z.-tritici and in an earlier publication (Stapley and McDonald, 2023). In brief, RADseq reads were trimmed of adaptors and low-quality sequences using trimmomatic (v0.35). The RADseq reads were mapped to a reference genome using bwa mem (v0.7.17). Reads from progeny of the 3D7×3D1 cross were mapped to the 3D7 reference genome and progeny from the 1A5×1E4 cross were mapped to the 1A5 reference genome. Variant calling was done using the GATK Germline Short Variant Discovery pipeline, following their Best Practice recommendations (https://gatk.broadinstitute.org/hc/en-us/sections/360007226651-Best-Practices-Workflows). After this we applied the following filters: only biallelic SNPs, parents had alternative alleles, <50% missing genotypes per marker, <50% missing genotypes per individual, mean read depth >3 and <30, depth quality (QD)>5, mapping quality >40, minor allele frequency >0.02 and <0.80, and SNPs in regions where the SNP density is >3 in 10bp were removed. Linkage maps built using these SNPs were described previously (Stapley and McDonald, 2023). After filtering and map building, 63181 SNPs were available for analysing the 3D7×3D1 cross and 32806 SNPs were available for the 1A5×1E4 cross. A summary of the two linkage maps is provided in Table S5 and the complete maps are available at (https://github.com/jessstapley/QTL-mapping-Z.-tritici).

### Phenotyping

We compared *in vitro* colony growth and melanisation at 18°C (control/benign environment) to that under cold stress (10°C) and heat stress (27°C). The optimal temperature for *Z. tritici* growing in vitro is 20.3°C (Tmin = 4.9, Tmax = 33.5) (Chaloner et al., 2021). Only non-clonal offspring strains passing all filters were phenotyped (for 1A5×1E4 n= 259 strains; for 3D7×3D1 n=265 strains). The protocols for isolate recovery from −80°C storage, growth in vitro and the measurements of colony size and colony melanisation were described previously (Zhong et al., 2021). In brief, Petri dishes containing potato dextrose agar (PDA) were inoculated with 200μl of a spore solution (concentration 200 spores/ml) and grown at 18°C for 8 and 12 days. Three replicates were grown at all temperatures. Two traits were measured; colony area and colony grey value, using automated image analysis as described previously (Lendenmann et al., 2014; Zhong et al., 2021). Grey value was measured on a scale of 0-255, where darker, more melanised colonies have lower values (0 = black, 255 = white). Mean colony area and grey value were calculated from measurements of multiple colonies on each plate and then the mean was calculated across the three replicate plates to obtain a final mean colony area and mean grey value for each strain in the two environments. Colony area was converted to a radial size measure by dividing by π and taking the square root of this value.

As colony growth varied with temperature and cross, it was not possible to make all measurements at the same days post inoculation (dpi) in both crosses. At 10°C colonies were measured at 12 and 15 dpi, at 18°C (control) colonies were measured at 8 and 12 dpi, and at 27°C colonies were measured at 12 and 15 dpi in the 1A5×1E4 cross and at 8 and 12 dpi in the 3D1×3D7 cross. Thus, it was not possible to directly compare colony measurements for the same colony age at 10°C and 18°C.

For the analysis we used three different types of measurements; measurements at a single time point (dpi) representing a specific colony age: including the colony radius and grey value at 8, 12 and 15 dpi; daily rate measurements: the change in colony radius or grey value between ages: 8-12 dpi or 12-15 dpi (e.g. the difference in colony radius between 8 and 12 dpi/difference in days); and tolerance measurements: the ratio of a measurement (colony radius, grey value, growth rate or melanisation rate) between a stressful temperature environment and the control environment (e.g. radius at 8 dpi at 27°C /radius at 8 dpi in 18°C environment, or growth rate at 27°C /growth rate at 18°C). For tolerance measurements in growth rate and melanisation rate, only heat tolerance (27°C compared to 18°C) could be calculated in the 3D7×3D1 cross, because growth rate and melanisation rate were measured over the same interval (8-12dpi) at 27°C and 18°C. The same tolerance values are not available for the 1A5×1E4 cross, or for colonies grown at 10°C, because the colonies were measured at different times between 18°C and 10°C. For the tolerance measurements, values >1 indicate the isolate is more tolerant to temperature stress (grew faster under temperature stress), and values <1 indicate that the isolate is more sensitive to temperature stress. In addition, at 12 dpi and 27°C there was no colony growth of the 1E4 parent and 43% of the offspring in the 1A5×1E4 cross did not grow. So, we calculated a binary trait value – colony growth or no colony growth – and searched for QTL associated with this binary trait using a binary model. In total there are 30 traits in the 3D7×3D1 cross and 23 traits in the 1A5×1E4 cross (Table 1).

### QTL mapping

QTL mapping was performed using the R (v 3.6.0) package ‘qtl2’ (v2_0.24) as described in detail on github (https://github.com/jessstapley/QTL-mapping-Z.-tritici). We scanned the genome for a single QTL per chromosome with the ‘*scan1*’ function using a linear mixed effect model. For continuous traits we fitted the kinship matrix in the model to control for the relatedness of individuals (i.e. we included a random polygenic effect). Models that take into account the genetic covariance between individuals can reduce the false discovery rate in QTL scans and outperform models that do not include this information in the model (Malosetti et al., 2011). For the binary trait we fitted a binary model, without the kinship matrix as this is not possible with this R package. The significance threshold for a QTL peak was determined by 1000 permutation tests and we calculated a Bayes Credible Interval (95%) to identify the interval size around the QTL peak. Overlapping QTLs were grouped to identify regions uniquely associated with traits in stressful and benign temperatures. QTLs were placed in the same group if the peak marker of one QTL interval overlapped with the 95% Bayes credible interval of another QTL.

### Identifying putative candidate genes in unique temperature stress QTL

A QTL uniquely associated with temperature stress is one where the peak marker for a QTL associated with temperature stress (10°C or 27°C) did not fall within the 95% credible interval of the QTL interval associated with a trait measured at 18°C. We focus on these unique temperature stress QTL as these provide the best opportunity to find genes specifically related to temperature response. Within each unique temperature stress QTL interval, we gathered information about each gene in both parents to identify the most likely candidate genes associated with temperature stress. This gene information included; the predicted gene function -paying particular attention to genes with known roles in temperature stress response (Temp-genes, Table S1), sequence (gene and amino acid) variation between the parents, expression differences between parents *in vitro* and *in vivo*, GO enrichment, and if the gene contained SNPs that were previously found to be associated with climate variables in a previous global environmental GWAS (Feurtey et al., 2023).

### Gene orthologues, function and variation in parents

We used custom R scripts to extract the list of genes within the interval from the annotation (gff) files of each reference genome. The putative encoded function of each gene was determined in previous analyses (for 1A5 see (Plissonneau et al., 2018), for 3D7 see (Plissonneau et al., 2016)). The annotations were obtained using Interproscan (https://www.ebi.ac.uk/interpro/) against multiple protein databases and then screened using multiple methods (i.e. signalP https://services.healthtech.dtu.dk/service.php?SignalP-5.0, Phobius https://phobius.sbc.su.se/) to determine if the encoded proteins contained likely signal peptides or secreted domains. The putative effect of a SNP in the coding sequence was determined using SNPeff (http://pcingola.github.io/SnpEff/). We identified the orthologous genes in the non-reference parent (3D7, 1E4) using an analysis performed across the genomes of 19 reference strains (Badet et al., 2020). We compared the DNA sequence and amino acid similarity between parents. DNA sequences were compared with BLAST and amino acid sequences were aligned and compared using R:protr (Xiao et al., 2015).

### RNAseq data analysis

We analysed RNAseq data that was previously created for the four parental strains during infection of the susceptible wheat cultivar Drifter (Palma-Guerrero et al., 2017) as well as cultures grown *in vitro* on two types of liquid media (yeast sucrose broth (YSB): yeast extract 10 g/L and sucrose 10 g/L, pH 6.8; and carbon-depleted minimal medium (MM), pH 5.8 (Francisco et al., 2019)). The analysis was described previously (Stapley and McDonald, 2023). The raw sequence reads were downloaded from the Short Read Archive (SRA, https://www.ncbi.nlm.nih.gov/sra, Bioproject: *invitro* SRP152081, *invivo* SRP077418). All code and input/output files are available on github (https://github.com/jessstapley/QTL-mapping-Z.-tritici/). In brief, reads were trimmed and then mapped to one of two reference genomes (reads from 3D1 and 3D7 were mapped to the 3D7 reference genome while reads from 1E4 and 1A5 were mapped to the 1A5 reference genome). Then the number of reads mapping to each gene was counted using R::Rsubread (Liao et al., 2019) and we tested for differential gene expression between strains, 3D7 versus 3D1 and 1A5 versus 1E4, using R::EdgeR (Robinson et al., 2010).

### GO Annotation

We took all the genes within the unique temperature stress QTL intervals and grouped them according to their trait/temperature combination – e.g. colony size related traits at 10°C, or all melanisation related traits at 27°C. GO enrichment analysis was performed following this tutorial for non-model species (https://archetypalecology.wordpress.com/2021/01/27/how-to-perform-kegg-and-go-enrichment-analysis-of-non-model-species-using-r/). GO annotations were retrieved from the annotation files described above and we used the R package ‘topGo’ (Alexa and Rahnenfuhrer, 2023) to perform a Fischer test on the genes within each QTL interval following the associated guidelines. All three ontologies – Biological Process (BP), Cellular Component (CC) and Molecular Function (MF) were analysed. The background set for GO analysis was the entire gene/transcript set for each reference genome (3D7 and 1A5). KEGG pathway analysis was not performed because K numbers can be found for only 34% and 49% of transcripts for 3D7 and 1A5 respectively.

### Heat shock protein enrichment in unique temperature QTL

Heat shock proteins (HSP) are numerous (44 annotated in our reference genomes) and widely distributed across all core chromosomes in the genome. We tested if the number of HSP genes within a unique temperature QTL interval (*Obs_HSP_*) was more than expected by chance using a simulation approach. For each unique QTL interval we created 10,000 random QTL intervals of the same size and on the same chromosome. We counted the number of HSP genes in these random intervals, created a frequency distribution and calculated the probability that a random QTL was equal to the number we observed (*Obs_HSP_* /10,000). If the probability was less than 0.05 then we concluded that the observed value was greater than expected by chance.

### Overlap with GWAS SNPs associated with climate

We investigated the overlap between genes within our intervals and those found in previous GWAS analyses. Earlier GWAS analyses used different reference genome annotations, hence the approach we used to find overlaps between the TSQTL and the GWAS significant SNPs differed according to the comparison. For the trait-based GWAS (traitGWAS) (Dutta et al., 2023), many traits were measured, but we only consider associations with growth and melanisation traits measured at 15°C, or relative growth/melanisation between 15°C and 22°C. The SNPs and their genomic positions were reported in five refence genomes (IPO323, 3D7, 3D1, 1A5, 1E4), so we could directly compare gene IDs from our QTL intervals to the traitGWAS results. For the other two GWAS analyses (Feurtey et al., 2023; Miñana-Posada et al., 2024), IPO323 was the reference genome, thus we had to convert IPO323 gene names to the orthologous positions in our parental reference genomes. In the genotype-by-environment GWAS (geaGWAS), we used the list of SNPs having significant associations with bioclimatic variables associated with temperature and rainfall (for details see (Feurtey et al., 2023)). For the thermal adaptation GWAS (adaptGWAS) (Miñana-Posada et al., 2024) we used a list of SNPs with significant associations with traits measured at low temperatures only (12°C, 15°C). We used custom R scripts to find overlaps between these genes/SNPs and our unique temperature stress QTL intervals (TQTL) and the geaGWAS and adaptGWAS.

## CRediT authorship contribution statement

Jessica Stapley: Formal analysis, Data curation, Writing - Original Draft, Writing - Review & Editing, Resources, Data curation. Ziming Zhong: Methodology, Investigation, Data Curation. Bruce A. McDonald: Conceptualization, Methodology, Writing - Review & Editing, Supervision, Project administration, Funding acquisition.

## Data availability

All data is available at ETH Data repositories: for SNP data DOI: 10.3929/ethz-b-000550424 and for phenotype data DOI: XXXX. All code and details of the analysis are available at (https://github.com/jessstapley/QTL-mapping-Z.-tritici).

## Acknowledgments

We thank Tiziana Valeria Vonlanthen, Bethan Turnbull, Susanne Dora, Sarah Furler, Alexandra Waltenspühl, and Jasmin Wiedmer for collecting the phenotypic data and Javier Palma-Guerrero for supervision. We thank Marcello Zala for assistance in the laboratory and the Genetic Diversity Center (GDC) of Zurich for bioinformatic support. This research was supported by the Swiss National Science Foundation (31003A_155955 granted to BAM).

